# 96-Well Oxygen Control Using a 3D-Printed Device

**DOI:** 10.1101/2020.11.12.379966

**Authors:** Adam Szmelter, Jason Jacob, David Eddington

**Author notes:** **Corresponding Author**, David Eddington.

## Abstract

Oxygen concentration varies tremendously within the body and has proven to be a critical variable in cell differentiation, proliferation, and drug metabolism among many other physiological processes. Currently, researchers study the gas’s role in biology using low-throughput gas-control incubators or hypoxia chambers in which all cells in a vessel are exposed to a single oxygen concentration. Here, we introduce a device which can simultaneously deliver 12 unique oxygen concentrations to cells in a 96-well plate and seamlessly integrate into biomedical research workflows. The device inserts into 96-well plates and delivers gas to the headspace thus avoiding undesirable contact with media. This simple approach isolates each well using gas-tight pressure resistant gaskets effectively creating 96 “mini-incubators”. Each of the twelve columns of the plate is supplied by a distinct oxygen concentration from a gas-mixing gradient generator supplied by two feed gases. The wells within each column are then supplied by an equal flow-splitting distribution network. Using equal feed flow rates, concentrations ranging from 0.6% to 20.5% were generated within a single plate. A549 lung carcinoma cells were then used to show that O_2_ levels below 9% caused a stepwise increase in cell death for cells treated with the hypoxia-activated anti-cancer drug Tirapirizamine (TPZ). Additionally, the 96-well plate was further leveraged to simultaneously test multiple TPZ concentrations over an oxygen gradient and generate a 3D dose response landscape. The results presented here show how microfluidic technologies can be integrated into, rather than replace, ubiquitous biomedical labware allowing for increased throughput oxygen studies.

## INTRODUCTION

Inhaled oxygen enters the body at an atmospheric concentration of 21% and is transported by red blood cells where its consumption and diffusion vary tremendously from organ to organ, and even within individual tissues, resulting in a wide range of physiologic O_2_ concentrations^1^. Despite this, cells are most often grown in standard cell culture incubators at 21% O_2_ and 5% CO_2_. Additionally, tissue oxygenation is severely altered in pathological conditions such as, but not limited to, cancer, diabetes, stroke, and heart disease in which low O_2_ tensions result in hypoxia. This phenomenon is most actively studied in the field of tumour biology in which O_2_ acts as a “Janus molecule” resulting in both restrained proliferation and necrosis as well as an aggressive phenotype and resistance to chemotherapeutic agents^2^. Recently, several hypoxia-activated prodrugs (HAPs) have shown promise in treating various cancers by selectively targeting areas of low oxygen concentration within a tumor. However, none have yet to be made available for clinical use. A common mode of failure is off-target action in normal tissues that experience physiologic hypoxia; for example, the HAP, Tirapirizamine (TPZ), caused hypoxia-dependent retinal toxicity in mice^3^. Current studies are performed in hypoxia chambers or O_2_ control incubators which allow only a single oxygen concentration to be assayed at a time thus limiting the throughput and increasing time and expense. A more high-throughput method is needed to simultaneously screen the effectiveness of the drug at several oxygen concentrations to predict off-target toxic effects in-vivo thus saving time, resources, and potentially lives in animal and human testing.

Polydimethylsiloxane (PDMS)-based microfluidic devices have been investigated as a means for improved control over hypoxia chambers or incubators by our lab and others^4^. Due to its high gas permeability and direct contact with cell media, PDMS-based devices boast impressive control over oxygen concentration and can deliver precise spatial gradients^5,6^ with rapid equilibration—even on the order of seconds^7^. While powerful, these devices have been limited to small studies as they do not integrate with high-throughput equipment such as plate readers and require specialized fabrication processes. For this reason, our laboratory and others have developed custom PDMS-based devices that integrate with multiwell plates to allow for more high-throughput studies^8,9,10^. Despite this, no device has enjoyed mainstream use most likely due to the non-standard nature of PDMS in biomedical research. When in contact with cell culture media, PDMS has been shown to absorb and release small molecules^11^ such as drugs, growth factors, hormones, and signalling molecules thus reducing effective concentration^12^ and inhibiting normal cellular function^13^. Additionally, it may leach un-crosslinked oligomers^13^ and cause increased evaporation and bubble formation^14^. These issues prevent re-use of devices requiring fabrication of a new device for each experiment. Furthermore, variations in cross-linker ratio, curing time, and surface treatment may change properties such as stiffness and surface chemistry resulting in poor initial attachment and growth of certain cell types^15^. This may yield inconsistent results depending on device preparation methods.

The 96-well plate is a ubiquitous, essential tool within the pharmaceutical and biomedical research community. It consists of a flat-plate with a 8 x 12 array of “wells” that serve as small test tubes that are used in almost every application of life science for a wide variety of purposes ranging from growing cells and drug screening to optical detection, storage, and reaction mixing. Microplates and related products are a $1.5B/year industry^16^ with an enormous catalog of dedicated specialized plate-specific instruments and accessories such as plate readers, washers, and imaging systems. Rather than replace the current industry standard, an increasing number of laboratories are creating microfluidic devices designed to fit with existing multiwell plates ^17–20^. This demand is evidenced by microfluidic design and fabrication houses, such as Microfluidic ChipShop, who now offer multiwell plate integration in their catalogs. Previously, our laboratory has designed gas control inserts for both 6 and 24-well plates ^21,22^, however, they required special cylinders for each gas condition. Additionally, these devices required submersion in cell media making media changes messy and prone to contamination. Biologist collaborators who used the devices observed negative effects on cell growth from the undesirable contact of PDMS or 3D-printed resin with the growth medium. The device presented in this work (Fig. 1) overcomes these problems by seamlessly integrating into 96-well plates without contacting growth media to feasibly bridge the gap between individual university laboratory and more mainstream use.

**Figure 1:**
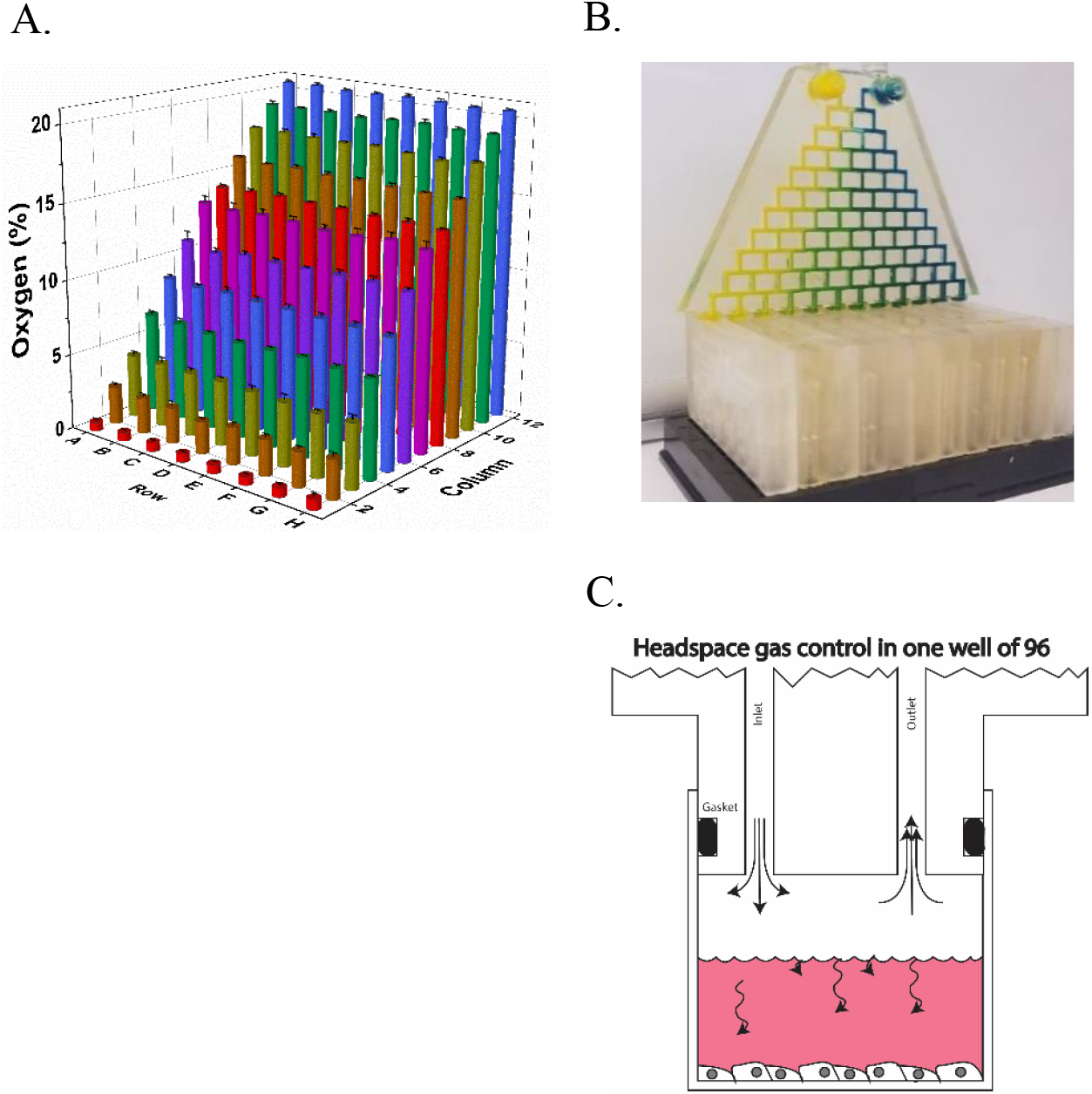
3D-printed 96-well gas delivery. A. Gas concentrations within each well of the device. B. Photograph of assembled device. Each column of the device receives a unique gas concentration supplied by the gradient generator. C. Schematic of headspace control. The flow to each column is equally split amongst the 8-wells through gas splitting distribution networks which deliver gas to the head-space.

## EXPERIMENTAL SECTION

### Device design and fabrication

Devices were designed in Solidworks and printed using a desktop stereolithographic 3D-printer (Form-3, Formlabs, USA) using the 25μm resolution setting. Devices were washed in the Formlabs Formwash resin removal device for 30 minutes using 70% (vol/vol) isopropanol (IPA) followed by flushing of internal channels using a syringe, also with 70% IPA. Next, the device and internal channels were dried with clean compressed air. Prints were then cured in the Formlabs FormCure for 1 hour at 60°C and let cool to room temperature before using. Buna-N O-rings (1mm thickness x 5.5mm OD) were manually placed on the pillars of distribution networks and the gradient generator. An important note is while it is simple to print a device, the time to print 12 distribution networks for an entire 96-well plate is approximately 144 hours. 3D printing is an excellent option for prototyping and open distribution but is not a practical solution for production at scale with current machines.

### COMSOL theoretical modeling

The designs for the gradient generator and distribution networks were evaluated using the finite element analysis modelling software COMSOL Multiphysics. For the gradient generator, a two-dimensional representation of the design was constructed, and gas mixing was evaluated using Laminar Flow and Transport of Diluted Species modules at 37 °C. The distribution network was imported as a 3D model created in Solidworks and flow rates were calculated according to the Laminar Flow module at 37 °C.

### Device assembly and operation

A distribution network was inserted into each column of the plate using gentle pressure. The gradient generator was connected to the twelve distribution networks such that each outlet of the gradient generator interfaced with each inlet of a single distribution network. The two feed gas lines, 21% O_2_/5% CO_2_ gas line and 0% O_2_/5% CO_2_ gas lines, both balanced with nitrogen, were connected to the two inlets of the gradient generator using Luer adaptors and flow rates were set to 100 SCCM using mass flow controllers (Alicat Scientific, MC-100 SCCM). Cheaper, regulated floating-ball mass flow meters may also be used, but tend to fluctuate at lower flow rates. Each of the two feed gases were humidified by bubbling through an Erlenmeyer flask containing 200 ml of sterilized DI water using stainless-steel carbonation stones with a 0.5μm pore size. Gas is humidified directly before entering the device. A filter may be attached to the inlet ports, however, experiments performed with or without filters did not result in any contamination and filters were not found to influence humidity (data not shown). Devices were sterilized by O_2_ plasma for 30 seconds or by briefly submerging in 70% ethanol and allowing to air dry in a sterile environment. They may also be autoclaved if printed using Formlabs High-Temperature or Dental SG resin. Before cell culture experiments, devices were warmed to 37°C in an oven or cell culture incubator before connecting to plates and gas lines to avoid condensation of humidified gas within channels. Each well may be seeded and filled up to a maximum of 187 μl of media which greatly exceeds amounts typically used in mammalian cell culture.

### Device validation using fiberoptic oxygen sensors

A fiberoptic oxygen probe (FOSPOR-R, Ocean Optics) was used to measure the oxygen concentration within each well. The sensor has a resolution of 0.0005% to 0.02% across the range of 0-21% oxygen and an accuracy of .01%. A two-point calibration was preformed within Ocean Optics software (Neofox Viewer, Ocean Optics) using humidified 0% O_2_/5% CO_2_ and 21% O_2_/5% CO_2_. To access the bottom of the well without allowing gas to enter or exit, a rubber septum access port was drilled for each well. Briefly, a Dremel tool was used to drill a 2 mm diameter hole in the bottom of each well which was subsequently sealed with a 2mm thick natural gum rubber sheet using a cyanoacrylate adhesive. The sheet was punctured using a specialized rubber septum puncture adapter for the oxygen probe (Needle Puncture Apparatus, Ocean Optics) and oxygen measurements were performed in each well without introducing gas or allowing it to escape. The oxygen probe was positioned at the bottom of each well by puncturing the rubber septum with the needle puncture adapter and positioning the probe approximately 100um from the tip of the needle as has been validated by others^8^.

### Burst pressure measurement

Distribution networks with single and double O-rings were evaluated for leaks and burst pressure. Devices were inserted into 96-well plates and house compressed air was connected to the inlets and a 100 psi gage pressure sensor (40PC100G, Honeywell). Outlets were sealed using a cyanoacrylate adhesive creating a closed circuit. Pressure was increased gradually while checking for leaks using a leak detection fluid (Leak Detection Compound, Cantesco). Pressure was logged to DAQAMI data acquisition software using a data acquisition unit (USB-1608FS-Plus, Measurement Computing) until the device physically dislodged itself (burst) from the plate. Burst pressure was recorded as the peak pressure attained.

### Evaporation and relative humidity measurement

Each well of a 96-well plate was filled with 150 μl of media and incubated at 37°C using inlet feed rates of 100 SCCM of humidified gas for 2 days. Loss in volume in each well was measured by pipette aspiration and results were spot-checked by measuring the change in weight of the entire plate and dividing by the number of wells. Relative humidity was assessed within each well using a humidity probe (GSP-6, Elitech) after 2-hours of humidification from bubbling through a stainless-steel carbonation stone (.5 μm pore size), 0.18” ID polyurethane pneumatic tubing, or no humidification. The humidity probe was positioned at the bottom of each well through modified rubber septum access ports as described in performing oxygen measurements.

### Cell culture

A549 lung carcinoma cell lines were grown prior to experiments at 37°C at 21% O_2_ and 5% CO_2_ and routinely split at confluence at a ratio of 1:4. The growth medium consisted of DMEM containing 10% FBS, 1% Penicillin and Streptomycin, as well as 4.5 g/dl glucose.

### Hypoxia-activated drug testing

Each well of a 96-well plate was seeded with 14,000 A549 Human Lung Carcinoma cells in 65 μl of media 24 hours prior to starting the experiment. In each column of the plate, 4 wells were grown in 33 μM TPZ and 4 wells were grown as controls in non-modified media. Each column was exposed to a different oxygen concentration ranging from 20.5% in column 1 to 0.6% in column 12. Cell viability was assessed daily through the use the non-destructive cell viability dye, PrestoBlue High Sensitivity (Thermo Fisher Scientific) in a plate reader (Varioskan Flash, Thermo Fisher Scientific) after the oxygenation network was removed from the plate. Fluorescence was measured using a plate reader with monochromator settings of 560/590 excitation/emission. Fluorescence intensity was normalized as fold change from day 0 (pre-experiment, 24 hours after seeding) levels. Each plate contained 4 technical replicates for each condition and four experimental replicates were performed to ensure statistical significance. For dose response experiments, each column was exposed to a different oxygen concentration ranging from 20.5% - 0.6% as above, and two rows were assigned for each of the four dose conditions: 0 μM, 25 μM, 75 μM, and 100 μM resulting in two technical replicates per condition. The experiment was repeated on three separate occasions.

### Statistical analysis

Oxygen concentrations generated by the device were validated in triplicate in which each well was measured on three separate days to ensure statistical experimental variance. Evaporation, humidity and burst pressure measurements were also validated in triplicate. Bar graphs depict mean values and error bars represent standard deviation. For the hypoxia-activated drug testing experiments, each well was normalized as fold change from a day 0 value taken 24 hours after seeding. Graphs represent the average of technical replicate wells per plate averaged over four experiments and error bars represent standard deviation. The HAP experiment had four technical replicates wells per condition per plate and the dose response experiment included 2 technical replicates per condition per plate. Single factor ANOVA in Origin (OriginPro, OriginLab) was used to calculate p-values between conditions.

## RESULTS

### Principle of device operation

An assembled device (Fig 1B) consists of a gradient generator (Fig. 2A) which mixes two gases over 10 generations into 12 outlets and twelve distribution networks (Fig 2C). Each outlet provides a unique gas concentration to a single column of the plate. To achieve equal flows within the wells of each column, a flow-splitting distribution network interfaces with the well-plate using a radial double O-ring design (Fig. 3B). This design is also used to connect the gradient generator to the twelve distribution networks. The insert device consists of a mirror image of the inlet and outlet delivery channels and constant flow drives gas into the well and out the outlet as shown in Figure 1B-C. Distribution networks are designed to split flow evenly into 8 equal resistance channels branching from a central hub (Fig. 2B,2D). COMSOL simulations confirm approximately equal distribution of flow rates (Fig. 2D).

**Figure 2:**
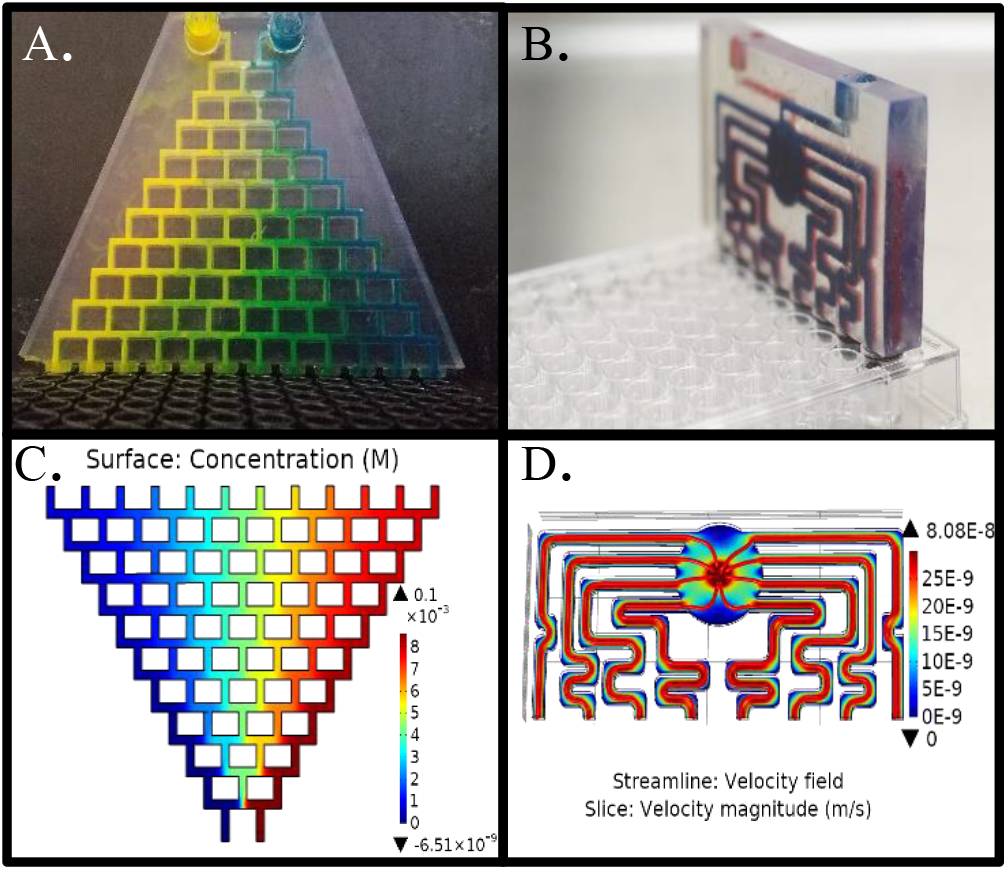
Device components: gradient generator and distribution networks. A. Photograph of the gradient generator injected with artificial coloring (food dye) to demonstrate mixing. B. Photograph of distribution network with red and blue dyes used to denote inlet and outlet channels. C. COMSOL simulation of gradient generator displaying 12 unique concentrations and D. COMSOL simulation of distribution network displaying equal distribution of flow rates (streamlines (red), surface velocity (colormap).

**Figure 3:**
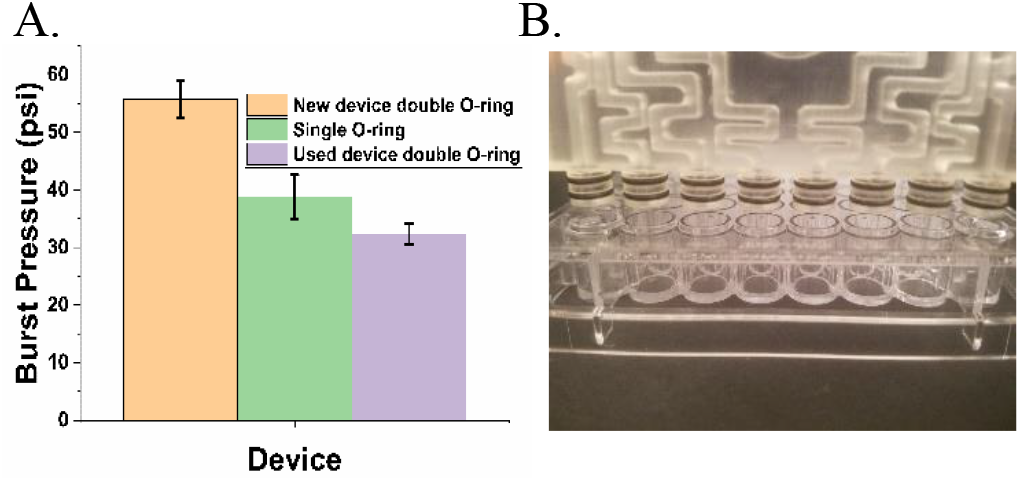
Burst pressure measurement of devices. A. Double O-ring devices were able to withstand higher burst pressures than single O-ring devices or old double O-ring devices (100+ cycles). B. Photograph of a single distribution network above a column of wells highlighting the radial double O-ring design.

### Oxygen concentrations

At 37°C, with equal flow rates of 100 ml/min, 21% O_2_/5% CO_2_ and 0% 0% O_2_/5% CO_2_ gas streams entered the device and produced 12 distinct oxygen concentrations ranging from 20.6% to 0.6% (Fig. 1A). Wells within each column were remarkably consistent as demonstrated by a low average standard deviation of 0.076%. Average oxygen concentrations in each row were: 0.6, 2.4, 4.2, 6.6, 8.7, 10.9, 13.2, 14.1, 15.8, 17.7, 19.2, and 20.5 percent.

### O-ring burst pressure testing

Radial double O-ring devices (Fig. 3B) sustained an average pressure of 56 psi before being displaced from the plate while single O-ring devices sustained pressures of 39 psi, and double O-ring devices subject to repeated use (100+ insertion and removal cycles over 6 months) could withstand 35 psi on average presumably due to gradual tearing of the O-ring from repeated use.

### Media evaporation and gas humidification

Media humidification resulted in an average loss of 1.29% volume per day. Results are reported in Figure 4A as volume lost, in μl, per day. Gas humidification with the carbonation stone produced an average relative humidity of 94% compared to 87% with tubing, and 12% seen with no humidification (Fig 4B). It is possible that the residual humidity seen with no humidification could be due to leakage through the rubber septum measurement port caused by repeated use.

**Figure 4:**
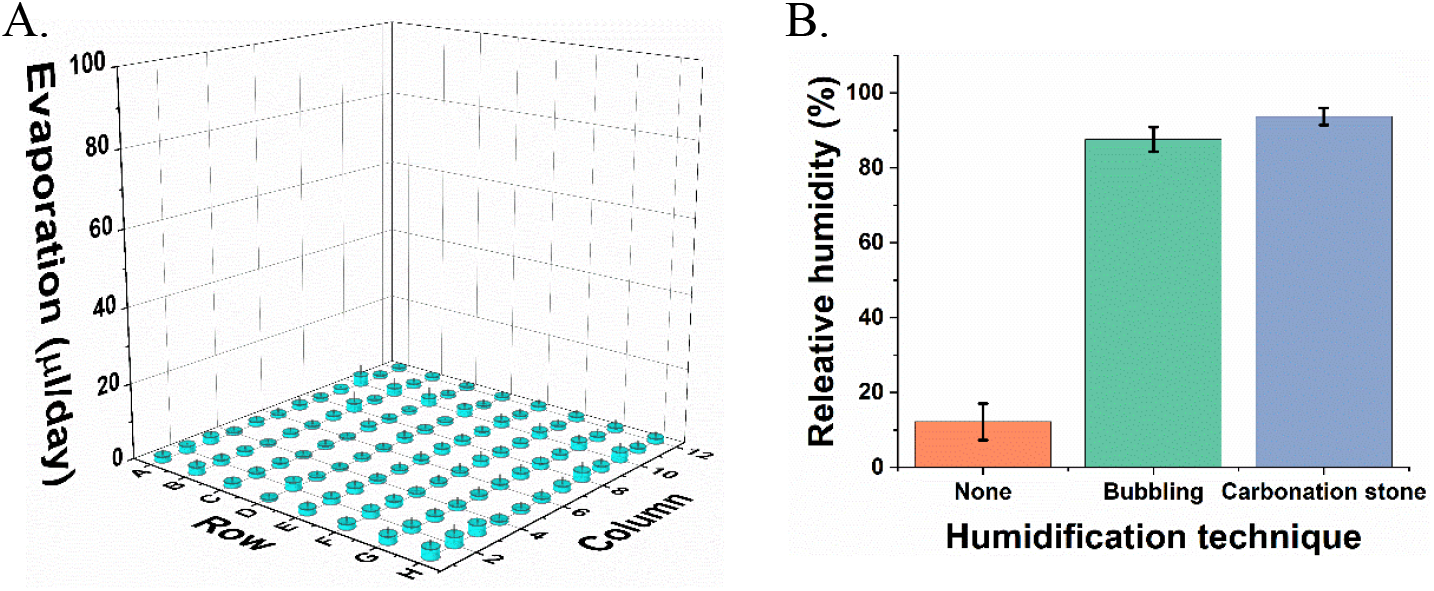
Evaporation and relative humidity measurement. A. Evaporation in μl/day from the each well of a 96-well plate. B. Relative humidity measurements within a well of a 96-well plate after humidification by carbonation stone, bubbling through tubing, or no humidification.

### Altering relative flow rates to change oxygen gradient range

Feed gas flow rates and concentrations were altered to narrow the range of oxygen concentrations delivered to the device (Fig 5). Mass flow controllers were used to increase the nitrogen stream flow rate relative to the 21% oxygen stream 5:1 (20 SCCM 21% O_2_5% CO_2_ and 100 SCCM 0% 0% O_2_5% CO_2_) narrowed the range of oxygen concentrations to 13.0% to 0% (Fig 5). By replacing the 21% O_2_ inlet gas with a 10% O_2_ line, and providing equal O_2_:N_2_ feed rates (100 SCCM 21% O_2_5% CO_2_ and 100 SCCM 0% O_2_5% CO), the concentration of gases was narrowed to 9.8-0.4% O_2_. By altering the relative flow rates of the two feed gases, or simply changing the feed gases themselves, it is possible to achieve any range of concentrations.

**Figure 5:**
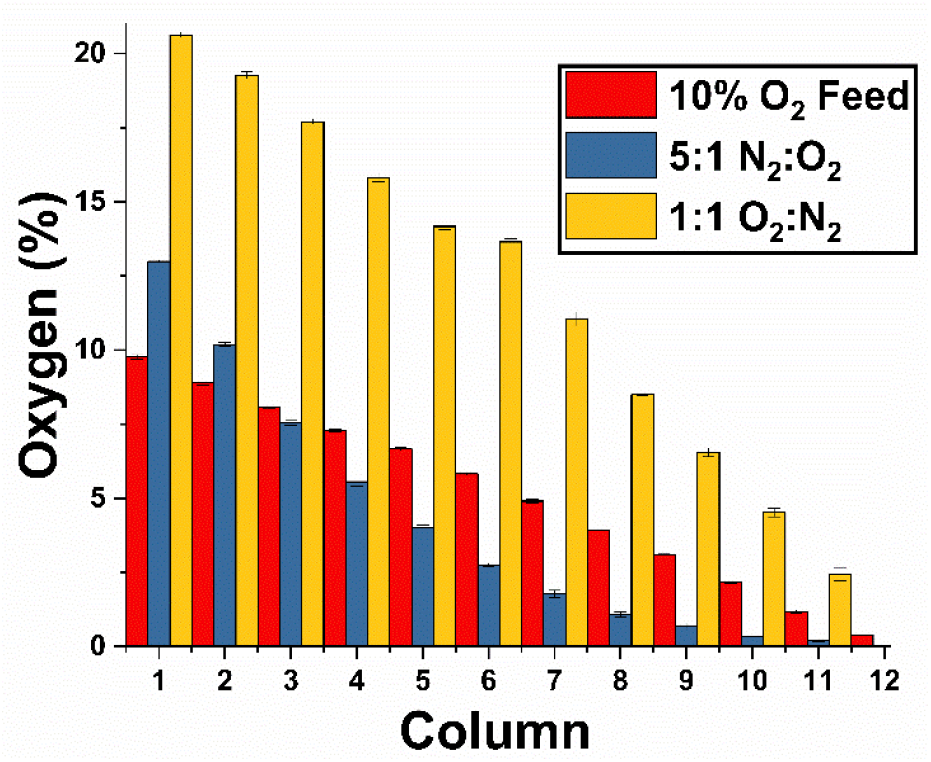
Altered oxygen concentration range through adjustment of relative feed flow rates. A ratio of 1:1 O_2_:N_2_ flow rates produced a gradient from 20.50% to 0.6% while a 1:5 ratio of O_2_:N_2_ flow rates produced a gradient from 13% to 0%. Supplying a 1:1 ratio of 10% O_2_ to 0% N_2_ produced a gradient from 9.7% to 0.38%.

### Hypoxia-activated drug efficacy testing

Oxygen concentrations ranging from 20.5% to 0.6% were delivered to cells treated with 33μM TPZ for 3 days. Viability was measured using the PrestoBlue assay at 0, 24, 48, and 72 hours. Each column of the plate received a distinct O_2_ level in which 4 of the wells were incubated with 33 μM TPZ while the remaining 4 wells were not given TPZ and treated as controls. A stepwise decrease in cell viability was seen in columns 8-12 corresponding to oxygen concentrations of 8.7% and below (Fig. 6). All columns showed a significant difference in survival compared to the no TPZ control, except columns 1 and 2, which were exposed to 20.5% and 19.2% O_2_, respectively. This may be due to insufficient activation of the drug at high oxygen tensions.

**Figure 6:**
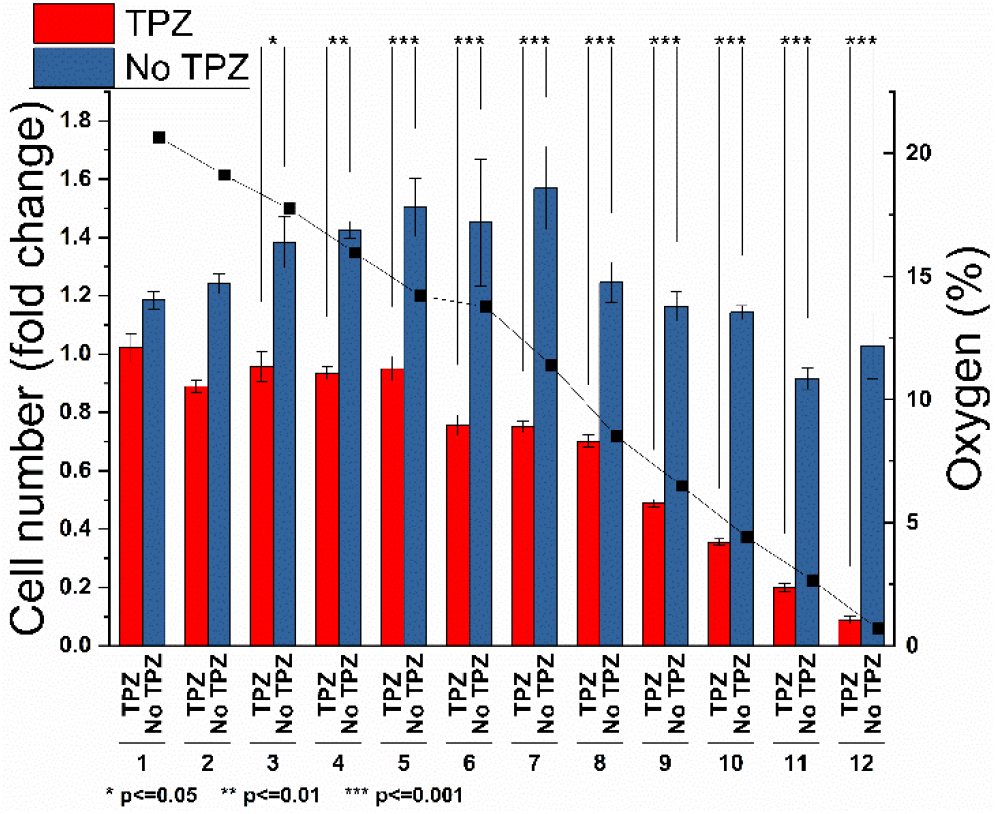
The hypoxia-activated prodrug, TPZ, is more effective at lower oxygen tensions. Twelve oxygen concentrations ranging from 20.5% to 0.6% were applied to a 96-well plate containing cells exposed to TPZ (red) or control (no TPZ, blue). Cells exposed to TPZ below 9% oxygen showed a linear decrease in survival. Asterisks denote significance between control and TPZ exposure groups.

A dose response experiment (Fig. 7) compared four doses of TPZ (0 μM, 25 μM, 75 μM, and 100 μM) across 12 oxygen conditions from 20.5% to 0.5% O_2_. Fold change in viability, compared to pre-experiment levels, was plotted as a threedimensional surface (Fig. 7A) with X/Y axis depicting oxygen concentration and TPZ dose and cell viability on the Z-axis. LD50 (median lethal dose) was plotted as a dotted line showing oxygen and TPZ concentrations that resulted in 50% cell death.

**Figure 7:**
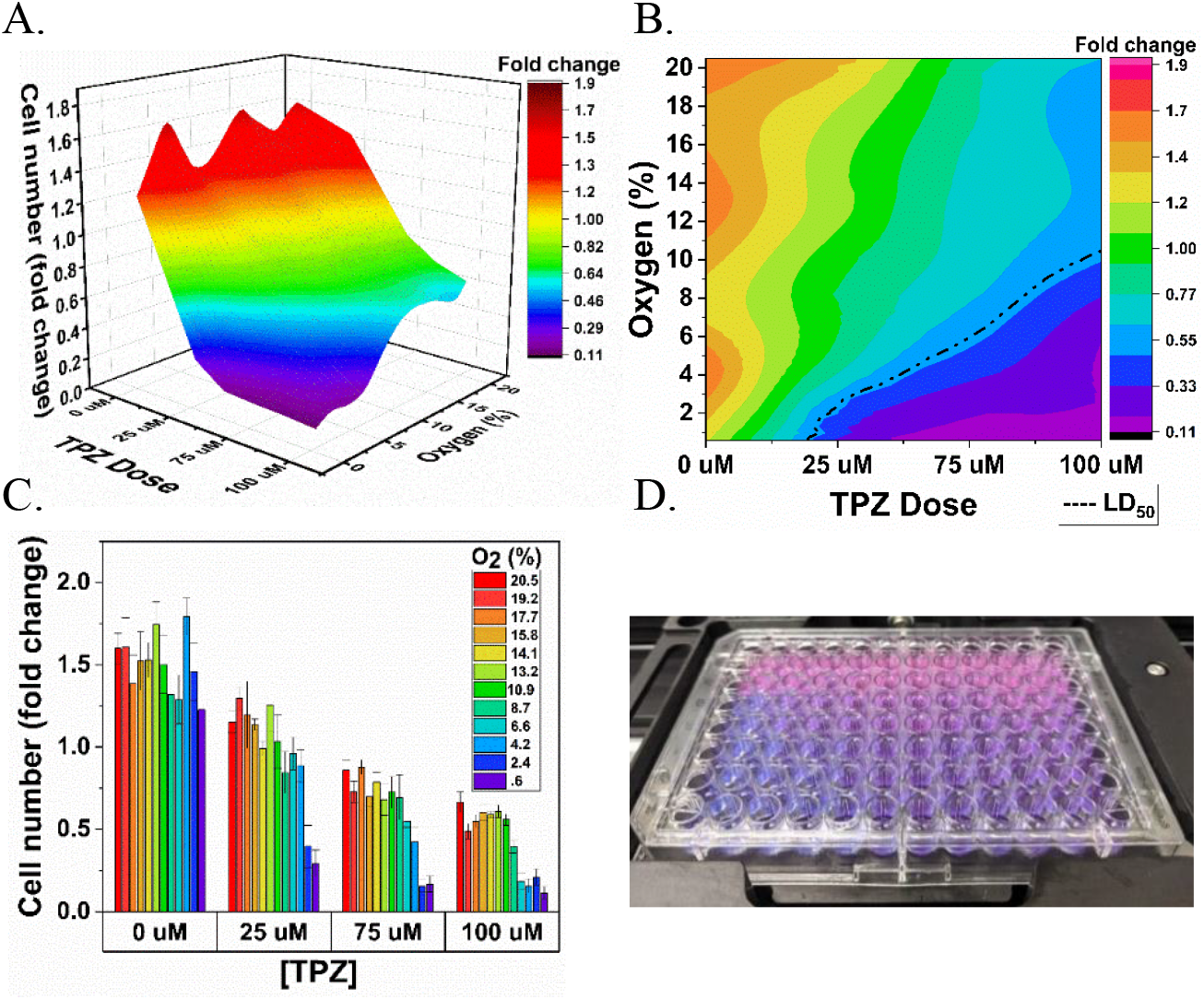
Oxygen and drug concentration landscapes on a single 96-well plate. A. Surface plot depicting cell viability (z-axis) under different oxygen and TPZ concentrations within a single plate. B. 2D plot depicting cell viability under different oxygen and TPZ concentrations. Oxygen concentration and drug dose combinations resulting in 50% survival are depicted by the dotted line. C. Bar plot depicting cell viability as a function of TPZ dose and oxygen concentration (colormap). D. Photograph of 96-well plate with PrestoBlue viability dye after two days of exposure.

## DISCUSSION

PDMS remains a major barrier preventing mainstream use of microfluidic devices for cell culture applications. Although its high oxygen permeability is useful for gas delivery, the material is widely considered to be “non-ideal” for cell contact, and its biocompatibility has been referred to as a “misnomer”^23^. Although treatments and surface modifications may improve certain aspects of biocompatibility, this may be too labor-intensive for most researchers who prefer out-of-the-box, standard tissue culture polystyrene^24^. Furthermore, variation in protocols may result in inconsistent results depending on the exact treatment protocol. For this reason, we have chosen to design a device that adapts to standard labware and can be easily integrated into biomedical workflows.

Our device dramatically increases the throughput of gas-control experiments compared to standard methods. Traditional experiments will use a single-vessel exposure system, such as an incubator or hypoxia chamber which supply a single oxygen concentration at a time. This approach can be extremely time consuming, especially for studies requiring 8 or more oxygen concentrations—such as is required for determining the potency of hypoxia-activated prodrugs^25^. By mixing two gases using a gradient generator and distributing them equally to the wells of each column of a 96-well plate using distribution networks, we produced a linear oxygen profile with 12 quantized concentrations, each with 8 replicates (Fig 1A). Finite element modelling using COMSOL Multiphysics was used to inform design and confirmed generation of 12 unique concentrations (Fig 2A) and equal splitting of flow rates to the wells of each column using the equal-resistance distribution networks (Fig 2B). While oxygen gradients produced in microfluidic devices can be robustly predicted and controlled through thorough modelling of variables such as fluid flow, reaction rates, and material properties^26,27^, isolating cells within a single well containing a single, quantized oxygen concentration has unique benefits for higher-throughput applications. Besides adapting into standard workflows, our method may be used to investigate O_2_ as a standalone variable, decoupled from factors such as fluid flow or interaction with nearby cells growing at different concentrations along the gradient. This may be particularly useful for deconstructing complex physiologic phenomena such as liver zonation, in which metabolism is spatially separated according to the availability of oxygen, hormones, and metabolites. Current microfluidic models of this phenomena have ascribed a varying, and sometimes contradictory, effect of O_2_ on enzymatic expression^28,29,30^. By isolating oxygen as a variable, our device may be used to elucidate the role of oxygen in complex physiologic or pathophysiologic phenomena.

To achieve accurate and reliable headspace oxygen concentrations, a robust leakproof, gas-tight seal is needed. This was achieved by incorporating radial double O-ring seals (Fig 3B) onto the pillars of the distribution networks and gradient generator. Burst pressure testing confirmed that distribution networks are leakproof up to 56 psi of pressure (Fig 3A). Interestingly, devices subjected to 100+ use cycles over 6 months showed lower pressure resistance (35 psi), presumably from O-ring tear or slight shrinkage of the 3D-printed device over time. Even with repeated use, the seal provides more than adequate protection against leaks. However, the strength of the seal presents additional opportunities in that the device may be adapted for studies in hydrostatic pressure biology in which simple valves connected to outlets could control venting and selectively allow pressure to build within the wells. Additionally, resistance to leaks at this high level of pressure allows for the safe delivery of toxic gases, gasotransmitters, and environmental pollutants which currently lack reliable in-vitro delivery methods^31^. Our system can be easily, and safely, adapted to these studies by placing the device within in a small oven set to 37°C within a chemical fume hood or by connecting tubing to vent outlet gas into a fume hood.

A major concern for all gas delivery devices is evaporation. This is particularly important for this device which relies on constant flow of gas above the cell culture media. We have addressed this problem by humidifying gas prior to delivery to the device as shown in Figure 4. Gas is bubbled through a stainless-steel microporous carbonation stone within a sealed Erlenmeyer flask to increase the contact surface area of the gas with the liquid. Our results show a slight improvement over evaporation within 96-well plates stored on the top shelf of a 95% humidity incubator^32^.

By delivering gas to the headspace, our device is convenient for biologists, however, this also subjects it to slower diffusion times compared to PDMS-based microfluidic devices. As within incubators or hypoxia chambers, gas dissolves from the headspace into the media according to Henry’s law, which governs the gas solubility in liquid, then to the bottom of each well according to Fick’s law, which relates diffusion time to distance (media height) and the concentration gradient. However, due to the low evaporation observed in our system, it is possible to use small media heights (<2mm) which minimize diffusion time and ensure oxygen flux through the media is more than sufficient to meet cell metabolic demmands^33^. While not ideal for fast-switching intermittent hypoxia experiments, this system is suitable for multi-day exposure experiments as shown in Fig. 6 and Fig 7 where we observed oxygen-dependent drug activity over two or three days. Additionally, the device can be placed on a heated microscope stage for long-term continuous cell imaging experiments. Here, cells may be observed during gas exposure which is not possible using end-point measurement techniques such as plate readers which require our device to be removed before each measurement.

It is important that a device be adjustable to the needs of individual users. While 0-21% O_2_ may be useful for investigating the broad toxicity of hypoxia-activated drugs within the body (Fig. 6-7), more targeted O_2_ levels are often required. In the body, oxygen concentrations vary dramatically, even within individual organs. As mentioned previously, tissues such as the liver have distinct metabolic zones delineated by oxygen concentration which can range from 4-14%^34^. By adjusting the relative flow rates of the feed gases within our system, or the concentration of the feed gases themselves, the device can achieve concentrations suitable for any physiological or pathophysiological system. This was demonstrated in Figure 5, where concentrations ranges were narrowed using 10% oxygen feed gas or by changing the feed ratio to 5:1 O_2_:N_2_. It is our hope that this flexibility will allow researchers studying different cell types, tissues, or diseases processes to tune oxygen concentrations specific to their needs.

To validate the biological usefulness of our device, we demonstrated a simple experiment in which a single 96-well plate was used to test the effectiveness of several concentrations of hypoxia-activated drug (TPZ) over 12 oxygen concentrations. First, we confirmed the stepwise increase in drug activity with decreasing oxygen concentration at a single drug dose (Fig. 6) by measuring cell viability over 3 days. This result confirms that a single plate can perform the work of 12 incubators in a small fraction of the space. For most laboratories, housing 12 incubators is neither space nor cost effective. However, without an entire room dedicated to incubators, this experiment would still take 36 days to complete with a traditional two incubator setup—this demonstrates an order in magnitude improvement in throughput. In this experiment, each column (supplied by a unique oxygen concentration) contained four replicates for drug treatment and four replicates for no-treatment controls. These technical replicates were useful for observing noise caused by the device or experimental protocol but did not take advantage of the power of the 96-well plate to test multiple variables simultaneously. To this end, we introduced TPZ dosage as a variable while still maintaining two technical replicates per condition. Four dosages were compared across the 12 oxygen concentrations to produce a map of cell viability under the two variables. Our results confirmed our hypothesis that increasing dosage and decreasing oxygen concentration result in increasing levels of cell death. A 2D plot allows for simple visualization of median lethal dose (LD50 values) at each oxygen concentration and shows the predictable response of decreasing with decreasing oxygen levels (Fig 7B, dotted line). Plotting the results in three dimensions produces a topographical landscape of cell viability under the varying oxygen and drug concentrations (Fig. 7A). Here, it is easier to visualize concentration combinations that result in steep changes in viability. This information is critical for determining safe concentrations that minimize off-target action in tissues which normally contain lower oxygen concentrations. A topographical plot shows the richness of information that can be contained within a single 96-well plate and, while not present in our data, can also be useful for identifying unusual behavior such as saddle-points and biphasic relationships.

Inexpensive desktop SLA 3D-printers are increasingly used by university laboratories and makerspaces to create tools, often microfluidic^35^, for scientific discovery^36^. Our device can be easily printed and used without the need for specialized photolithographic fabrication techniques or assembly. With the barrier to entry significantly lessened, it is our hope that this device will allow more laboratories to incorporate gas control in their research.

## CONCLUSIONS

Our contactless headspace delivery method avoids an undesirable interface with cell media and thus provides a convenient way to control gas while still integrating with standard 96-well plates and their associated workflows and instruments. By designing the device for production using a desktop SLA 3D-printer, we hope this will allow wider access to gas-control for laboratories who may now print the device and study the role of oxygen for a particular organ system, disease process, or drug study. Additionally, our closed-well system opens the door for potential use with biologically significant gases such as gasotransmitters and environmental pollutants which currently lack reliable methods for in-vitro delivery. The system is amenable to the oxygen concentrations seen within any organ system or disease process by adjusting the relative flow rates of the two feed gases, or the content of the feed gases themselves. This 3D-printed gas control insert is a powerful tool that offers flexibility, convenience, and the increased throughput needed for widespread use in the biomedical research community.

## AUTHOR INFORMATION

### Author Contributions

The manuscript was written through contributions of all authors. All authors have given approval to the final version of the manuscript.

## ACKNOWLEDGMENT

We gratefully acknowledge the financial support from the NIH under award number R21EB024003, the National Science Foundation and the industrial members of the Center for Advanced Design and Manufacturing of Integrated Microfluidics NSF I-UCRC award number IIP-1841473, and American Heart Association Predoctoral Fellowship Award #20PRE3508008 for Adam Szmelter. Additionally, we would like to thank Haseeb Syed for help with oxygen measurements and Bill Myers for taking photographs of the devices.

